# hTFtarget: a comprehensive database for regulations of human transcription factors and their targets

**DOI:** 10.1101/843656

**Authors:** Qiong Zhang, Wei Liu, Hong-Mei Zhang, Gui-Yan Xie, Ya-Ru Miao, Mengxuan Xia, An-Yuan Guo

## Abstract

Transcription factors (TFs) as key regulators play crucial roles in biological processes. The identification of TF-target regulatory relationships is a key step for revealing functions of TFs and their regulations on gene expression. The accumulated data of Chromatin immunoprecipitation sequencing (ChIP-Seq) provides great opportunities to discover the TF-target regulations across different conditions. In this study, we constructed a database named hTFtarget, which integrated huge human TF target resources (7,190 ChIP-Seq samples of 659 TFs and high confident TF binding sites of 699 TFs) and epigenetic modification information to predict accurate TF-target regulations. hTFtarget offers the following functions for users to explore TF-target regulations: 1) Browse or search general targets of a query TF across datasets; 2) Browse TF-target regulations for a query TF in a specific dataset or tissue; 3) Search potential TFs for a given target gene or ncRNA; 4) Investigate co-association between TFs in cell lines; 5) Explore potential co-regulations for given target genes or TFs; 6) Predict candidate TFBSs on given DNA sequences; 7) View ChIP-Seq peaks for different TFs and conditions in genome browser. hTFtarget provides a comprehensive, reliable and user-friendly resource for exploring human TF-target regulations, which will be very useful for a wide range of users in the TF and gene expression regulation community. hTFtarget is available at http://bioinfo.life.hust.edu.cn/hTFtarget.

## Introduction

Transcriptional regulation is a fundamental and vital process for general and condition-specific gene expression [1]. Transcription factors (TFs) are the key regulators involved in transcriptional regulation [2]. Most TFs recognize and bind to specific DNA sequences (named as TFBSs), leading to distinctly spatiotemporal expression patterns of target genes [3]. The disturbance of TF-target regulation can disrupt normal biological processes, which may result in severe damage and diseases [4]. Thus, the identification of TF-target regulation is an important issue for understanding transcriptional regulation underlying complex biological processes [5]. Moreover, investigating spatiotemporal TF-target regulations can provide comprehensive insights for dissecting the regulatory mechanisms of gene expression underlying cell status and diseases [6].

Chromatin immunoprecipitation sequencing (ChIP-Seq) technology provides convenience to systematically investigate target genes of a TF at the genome-wide level [7]. The accumulation of ChIP-Seq data poses great opportunities to integrate the interactions between TFs and their targets in different conditions [8]. Meanwhile, comprehensive utilization and visualization of TF-target data can offer systematic views for TF-target regulations. Although several resources, such as ReMap [9], CistromeDB [10], Factorbook [11], ChIPBase [12], and TRRUST [13], were conducted to display the relationships between mammalian TFs and their targets. Most of them do not provide convenient functions to directly investigate TF-target regulations. For example, ReMap and Factorbook present experimental designs of ChIP-Seq datasets and peak information instead of putative TF-target regulatory relationships, while TRRUST deposits TF-target regulatory networks using a sentence-based text mining approach. Meanwhile, integrating large-scale omics datasets for TFs (including TFBSs, target prediction, mRNA profiling, and epigenetic status of chromatin) can provide a comprehensive understanding for transcriptional regulatory repertoires [14].

In this study, we developed a database named hTFtarget, which curated a comprehensive repertoire of TF-target relationships for human and offered an almost one-stop solution for studies involved in TF-target regulation. Our hTFtarget has integrated thousands of ChIP-Seq datasets and epigenetic modification information to predict reliable TF-target regulations, and also provides valuable tools to predict potential co-association and co-regulation between TFs. hTFtarget can serve as a useful resource for researchers in the community of TF regulation and gene expression.

## Implementation

hTFtarget was implemented with HTML, JavaScript (https://www.javascript.com/), Python (https://www.python.org/), Flask (http://flask.pocoo.org/), Bootstrap (https://getbootstrap.com/), AngularJS (a model-view-controller frame for Javascript web service, https://angularjs.org), and WashU EpiGenome Browser [15]. MongoDB (a cross-platform document-oriented database engine, https://www.mongodb.com/) was used to store metadata information. hTFtarget was hosted on the Ubuntu Linux system (version 16.04) with the Apache HTTP Server to provide a stable and open service.

## Database content and usage

### Data collection and quality control

Non-redundant ChIP-Seq datasets of human TFs were curated from public databases, including NCBI GEO, NCBI SRA, and ENCODE [8]. For GEO and SRA databases, ChIP-Seq datasets were collected through the E-Utilities toolkit with filter criteria “gds or sra, human or Homo sapiens, ChIP-Seq or ChipSeq or ChIP sequencing, transcription factor or transcriptional factor or TF”. ChIP-Seq datasets from ENCODE database were enrolled using parameters “assay_term_name = ChIP-Seq, assembly = hg19/hg38, type = experiment, status = released, organism = Homo sapiens, target.investigated_as = transcription factor”. All datasets were manually curated to discard the non-TF and abnormal datasets, such as artificial TFs (mutated or fused), transcriptional co-factors, general TFII family members, and unclear descriptions for experimental designs. After the above procedures, FastQC (v0.11.5) was used for data quality control (QC) to obtain clean reads. Bowtie (v1.2.1) was employed to align clean reads to human reference genome GRCh38. Datasets with reads less than 5 M after the QC procedure or an alignment ratio less than 50% were discarded. Finally, 7,190 samples of 659 TFs from 569 conditions (399 cell line types, 129 tissues or cells, and 141 treatments) were kept for further analyses.

### Peak detection and motif discovery

Bam files of technically replicated samples from the same dataset were merged together, and the input or IGG samples were used as controls according to the experiment design (IGG samples were used as controls only for the absence of input samples). The peak calling procedure followed the protocol of MACS2 (v2.1.0) pipeline with the following parameters (q-value ≤ 0.01, fix-bimodal) [16]. Putative motifs of the TF in a dataset were identified using a similar method proposed by Wang et al. [17]. The detailed procedures were: 1) All of the peaks were ranked by the value of enrichment signal, and then the top 500 peaks (training set) were used for motif discovery using MEME-ChIP suite (v4.10.0) [18]; 2) The top five motifs (ranked by E-value, E-value ≤ 1e-5 and the “match sites” ≥ 100 in the top 500 peaks of step 1) were considered as confident ones, and the top 501–1000 peaks from the step 1 were served as the testing set to measure the power of the top five motifs. The same amount of random genomic regions of GRCh38 reference genome (non-peaks) with similar GC contents and sizes were selected as the control set. The FIMO (v4.10.0) [19] was used to scan both the testing and control sets, and then the numbers of recurrent motifs within the two sets were used to evaluate the significance of motifs for the TF (*t*-test with Bonferroni corrected p-value less than 0.01). Motifs with significant reoccurrence only in the test sets were considered as high-confident ones and used for peak filtration; 3) Peaks with a p-value ≤ 1e-5 and containing the above high-confident motif(s) were considered putative functional peaks and then used for further analyses.

### Identification of TF-target regulation

Based on the putative functional peaks, we implemented the beta-model to identify candidate TF-target regulation [20], in which the beta-model score was used to measure the potential power of peaks for the TF-target regulation.

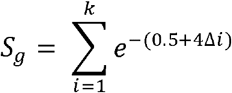

where *S_g_* is the beta-model score represented as the sum of the weighted scores of peaks nearby the TSS of gene *g*, and the parameter *k* is the number of TFBSs within the 50 kb distance from the TSS, while Δ*i* is the distance between the summit of peak *i* and the TSS (which was normalized to 50 kb; *e.g*., the value 1 represents a 50 kb distance, while 0.04 indicates 2 kb). For the extreme case, only one peak was detected within 50 kb, and the summit of the peak was exactly at 2 kb before the TSS, the beta-model score was 0.517. Thus, the value 0.517 was used as the cutoff for regulatory capacity. If the putative functional peak with a beta-model score ≥ 0.517 locates within 50 kb upstream of the TSS, we consider the TF-target regulation is reliable. Further analyses and functional modules in hTFtarget were based on reliable TF-target regulation.

Moreover, we integrated TF-target regulations from multiple datasets of each TF and the chromatin status of the upstream region of a target gene, to survey whether the TF-target relationship is general or condition-specific. First, for each TF, we combined all of the targets of the TF from multiple datasets together, and the upstream 50 kb region from the TSS of each target was divided to subunits with a bin length of 200 bp. Then we examined whether any of the subunits was labeled with activated epigenetic modifications (data source curated from Roadmap project, http://www.roadmapepigenomics.org/). Finally, if the TF-target regulation is supported by one peak with a beta-model score ≥ 0.517 in more than 30% datasets of the TF, and any of the subunits of the 50 kb upstream region for the target gene is labeled with an activated epigenetic modification status within at least 30% of the Roadmap epigenomic samples, we consider the TF-target regulation is a general case; otherwise it is a condition-specific one.

### TFs co-association detection

The co-association of TFs, which was predicted according to the method proposed by Mark et al. [21], indicates the probability of TFs co-binding to common genomic regions. Briefly, to investigate the co-associations between a given TF (focus-factor) and other TFs (partner-factors) in a specific condition (a cell line or experiment), we first collected the overlapping regions of peaks between the focus-factor and each partner-factor to obtain a systematic co-binding map. We then implemented a machine learning method, which combined the RuleFit3 algorithm and mutual information of the overlapping regions between the focus-factor and partner-factors, to calculate the quantitative relationship as the relative importance score between the focus-factor and each partner-factor. A higher relative importance score of two given TFs indicates a stronger and more confident co-association relationship between them.

### TFs co-regulation to target gene

Target gene co-regulated by more than two TFs was predicted based on the reliable TF-target regulations detected from curated ChIP-Seq datasets in hTFtarget. Briefly, for a query gene, the 50 kb region upstream from its TSS was served as a core area to predict the candidate co-regulation of TFs. TFs with high-confident peaks within the upstream regions of TSS of target gene and the beta-model score of each peak ≥ 0.517 were considered as putative co-regulators to the query gene.

### Sequence based TFBS prediction

For the prediction of candidate TFBS(s) on given sequence(s), we integrated the motif matrices of human from our hTFtarget database and other resources (HOCOMOCO [22], JASPAR [23], and TRANSFAC v2018 databases [24]). In total, we collected TFBSs for 864 human TFs, and the FIMO tool was employed for the TFBS discovery on given sequences.

### Data resource and web-interface features

hTFtarget deposited 3.2 million records of TF-target regulations for 659 TFs in 569 experimental conditions and integrated 2,737 high-confident motifs of 699 TFs from other databases to predict 3.5 million records of candidate TF-target regulations. Moreover, 20,320 motifs were curated in our hTFtarget database from ChIP-Seq data. Meanwhile, 408 TFs were found to possess co-association ability in 10 cell lines. Furthermore, hTFtarget has a user-friendly web interface to facilitate the search, browse, and download of comprehensive TF-target relationships in multiple experimental datasets, including putative TF-target regulations, ChIP-Seq peaks, epigenetic modification status of TFBSs and targets, co-regulation of TFs to targets and co-association of TFs. All of the data sources and functional modules in hTFtarget are shown in Figure 1. We also compared hTFtarget with other resources involved in TF-target regulation (Table 1). We found that our hTFtarget may be the most comprehensive database providing various functions for the exploration of TF-target regulations in human. Our hTFtarget is freely accessible at http://bioinfo.life.hust.edu.cn/hTFtarget/.

**Figure 1.**
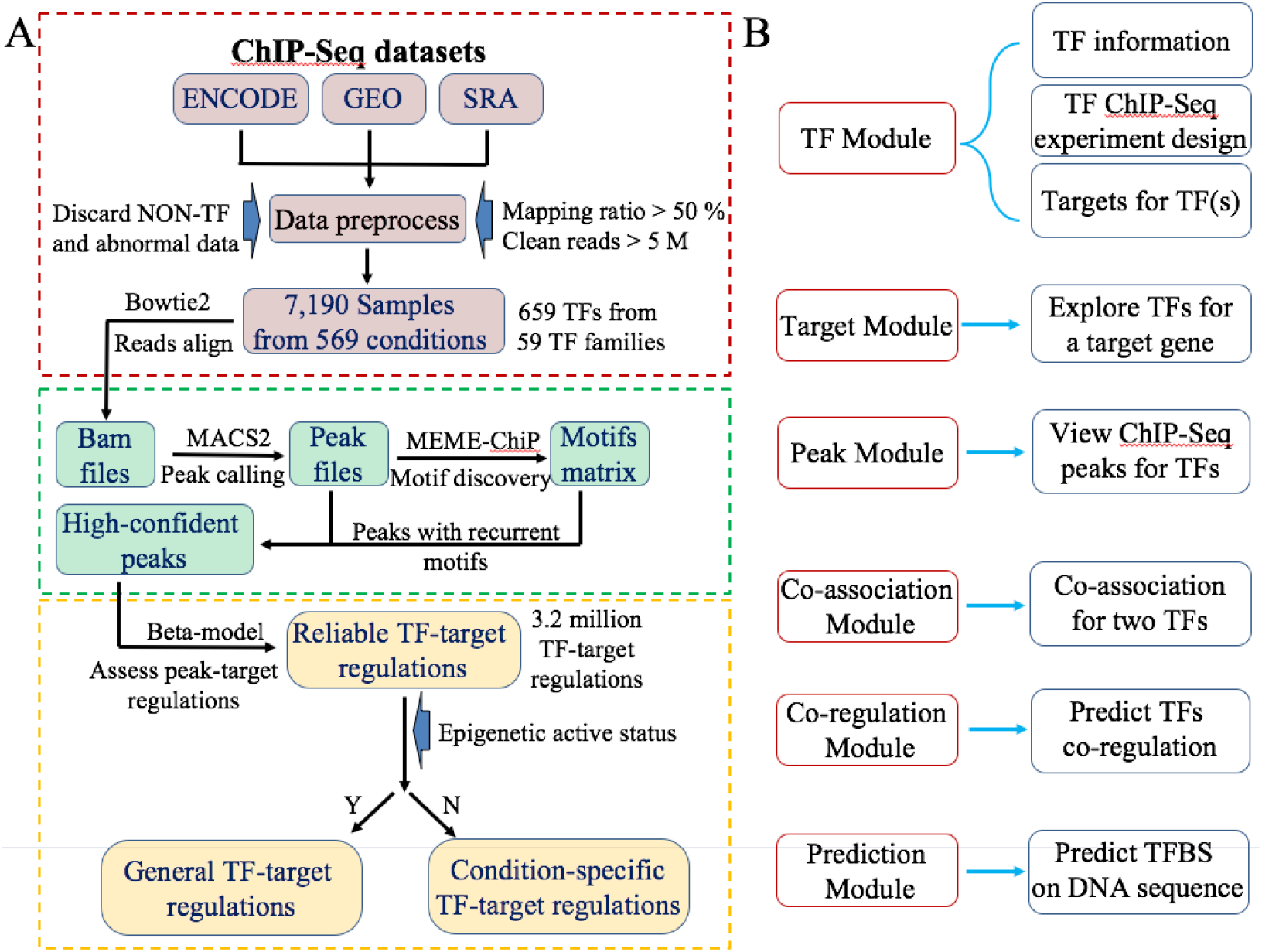
Overview of data resources and functional modules of hTFtarget. (A) The resource summary and workflow for the detection of TF-target regulations in hTFtarget. (B) The main functional modules of hTFtarget. TF: transcription factor; ChIP-Seq: Chromatin immunoprecipitation sequencing.

**Table 1.** Summary of well-known databases in TF-target regulation

|  |  | **hTFtarget** | **ReMap** | **CistromDB\*** | **Factorbook** | **TRRUST** | **ChIPBase** |
| --- | --- | --- | --- | --- | --- | --- | --- |
| **Data** | Source | Experiment | Experiment | Experiment | Experiment | Publication | Experiment |
|  | Technology | ChIP-Seq | ChIP-Seq | ChIP-Seq, DNase, ATAC | ChIP-Seq | Text mining | ChIP-Seq, DNase, ATAC |
|  | No. of TFs | 659 | 485 | 1,700 | 167 | 800 | 480 |
|  | No. of datasets | 7,190 | 2,829 | 13,976 | 837 | - | 2,498 |
|  | No. of cells or conditions | 399/170 | 346/0 | -/- | -/- | -/- | -/- |
|  | Species | Human | Human | Human and mouse | Human | Human and others | Human and others |
| **Function** | TFBS prediction | Y | N | N | N | N | N |
|  | TFs co-association | Y | N | N | N | N | N |
|  | TFs co-regulation | Y | N | N | N | N | N |
|  | Target search for TF | Y | N | Y | N | Y | Y |
|  | TF search for target | Y | N | Y | N | N | N |
|  | Peak view | Y | Y | Y | N | N | Y |
|  | Peak comparison | Y | N | Y | N | N | N |
|  | Epigenetic status | Y | Y | Y | Y | N | Y |

*Note:* \*CistromDB collected a large number of transcriptional co-factors, RNA polymerases, TFII family members and their compounds, which resulted in the extremely high number of repository of TFs and datasets. TF: transcription factor; TFBS: transcription factor binding site; ChIP-Seq: Chromatin immunoprecipitation sequencing; ATAC: Assay for Transposase-Accessible Chromatin

### Usage

First, hTFtarget provides a “Quick Search” function on the top right of each page to conveniently survey the TF-target regulation for users. Various inputs are acceptable, such as the gene name, the Ensembl gene ID, gene symbol and alias. Users can obtain basic information for the query TF or gene, related TF-target regulations, epigenetic modification status in the promoter region of target gene, and detailed evidence for the TF-target regulation in different conditions. For example, when input gene *ASCL2* on the “Quick Search” function (Figure 2), all records of *ASCL2* as a TF or a target gene are shown in the Figure 2A.

**Figure 2.**
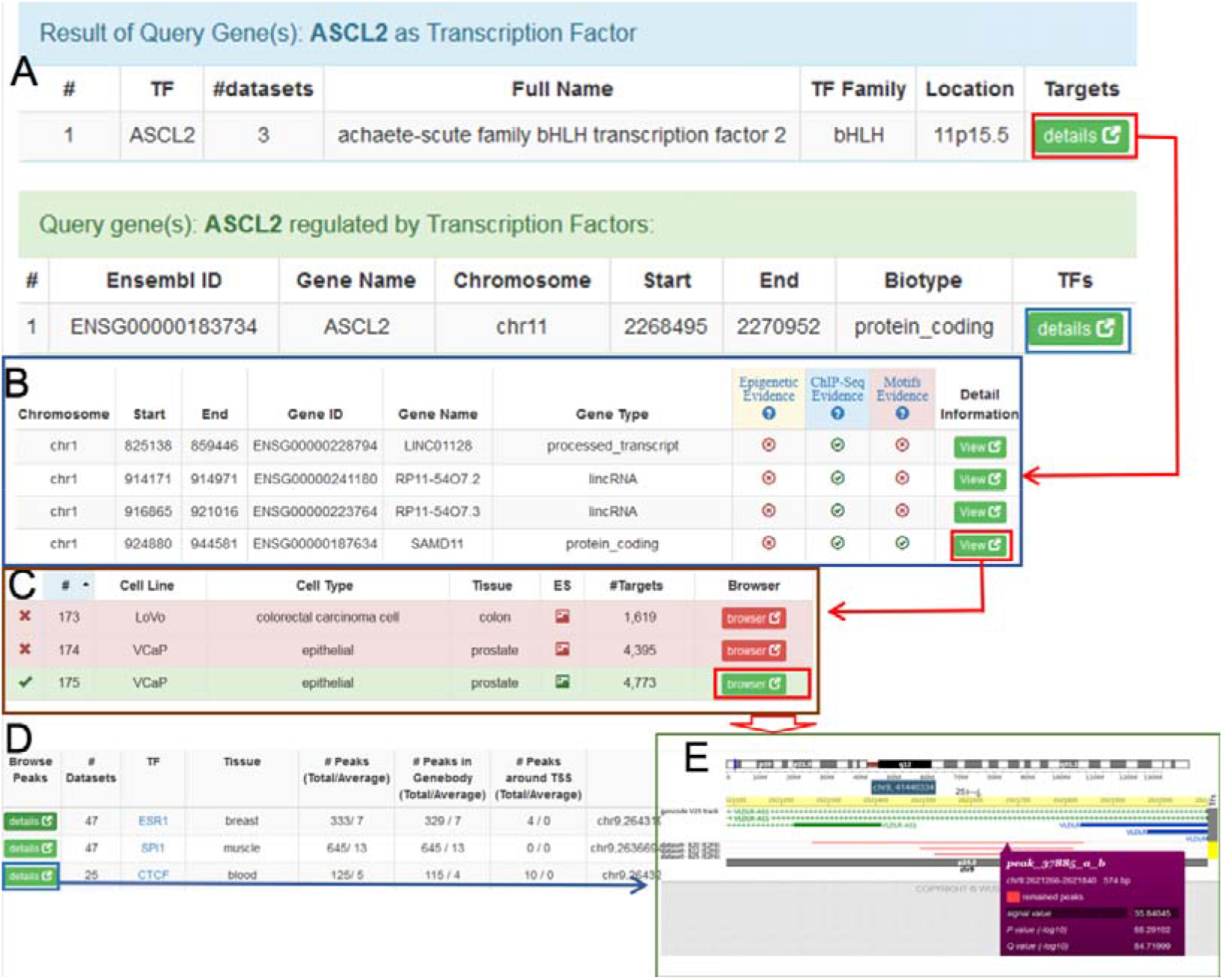
Snapshots of the quick search function in hTFtarget. (A) Results of a quick search function provide comprehensive views for TF-target regulations. (B) A partial screenshot of the results for the query gene as a TF after clicking the “details” icon. Each record represents a target regulated by the query TF. (C) Browsing of the condition-specific TF-target regulation in different experimental conditions or cell lines. (D) A part of the results for the query gene as a target. Each record indicates a TF-target regulation in a certain tissue. (E) The peak information of the query gene as a target.

On one hand, when the query gene *ASCL2* is a TF, the “details” icon below the term “Targets” (red box in Figure 2A) links to a page displaying its targets with three levels of evidence (Figure 2B). Additionally, users can filter target genes by supportive levels (“Epigenetic Evidence”, “ChIP-Seq Evidence” and “Motifs Evidence”) and browse the peak information through the “view” icon (Figure 2C). On the other hand, when the query gene *ASCL2* is a target, the “details” label underlying the term “TFs” (blue box in Figure 2A) shows all records about which TF could regulate gene *ASCL2* (Figure 2D). Users can browse the peak information for the TF-target regulation by through clicking the “details” icon (Figure 2D, Figure 2E).

Other than the quick search function, hTFtarget offers the following six modules for users to conveniently explore TF-target regulations (Figures 3 and 4).

1. TF module: provides details of TF-target relationships and ChIP-Seq experimental designs (Figure 3A). Users can explore the general (Figure 3A) or condition-specific TF-target regulations (Figure 3B), as well as browse the TF annotation (the bottom of Figure 3A) and the experimental design of the interested dataset (the bottom of Figure 3B). The “details” icons in Figure 3A enable users to browse the detailed information of TF-target regulations (Figure 3C), including the epigenetic status of the upstream region for the target gene and the visualized peak information in the genome browser (Figure 3D). Additionally, beyond surveying TF-target regulation in hTFtarget online, users can download the records as well.
2. Target module: helps users to investigate the TF(s) which regulate(s) the query gene. The function and the interface of this module are similar to Figure 2D and 2E.
3. Peak module: provides comprehensive information of peaks in a user-defined manner (Figure 4A). Users can browse the peaks of different TFs in the selected condition (*e.g*., the same cell line or tissue) or investigate TF-target regulations for the same TF in different conditions (*e.g*., the distinct spatiotemporal TF-target regulatory relationships in different tissues).
4. Co-regulation module: helps users to search the common targets which can be regulated by given TFs, or search common TFs which can regulate given genes (Figure 4B).
5. Co-association module: predicts and visualizes potential co-association TFs in collected cell lines. The co-association of two TFs indicates the probability of their combinatorial occupancies on the same genome regions. The human TFs have always shown distinct co-association relationships in combinatorial and context-specific patterns for gene regulation [21,25]. The relative importance score indicates the probability of co-association between two TFs, and the higher score means stronger co-association in the corresponding condition.
6. Prediction module: employs motif matrices to predict potential TFBSs on the user input sequence(s) (Figure 4C). The motif matrices were curated from TRANSFAC, JASPAR, HOCOMOCO, and hTFtarget databases.

**Figure 3.**
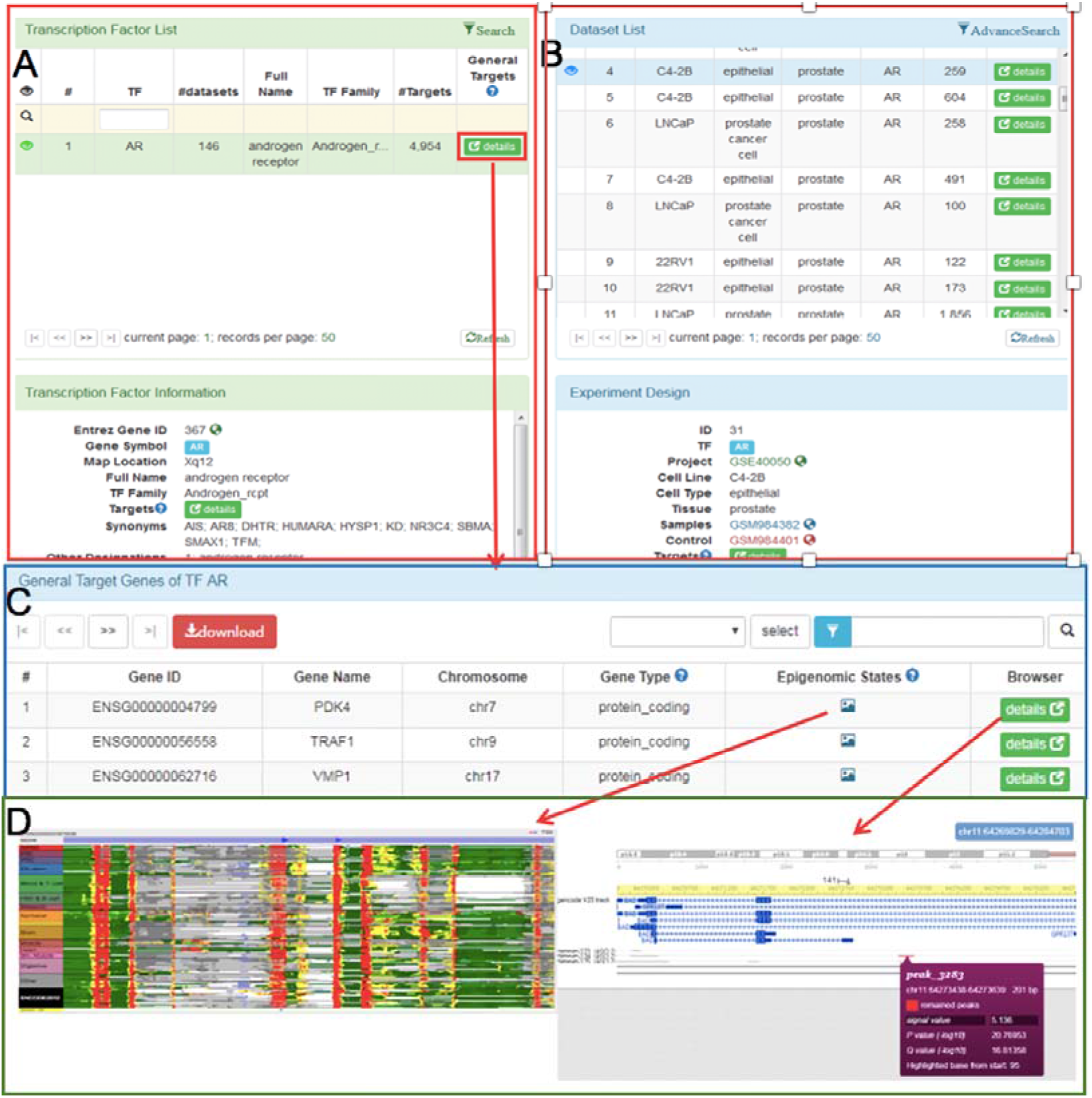
Views for the TF and target related modules in hTFtarget. (A) and (B) Basic information of the collected TFs. (C) General targets of the selected TF. (D) Epigenetic status and peak details within the flanking region of the target gene.

**Figure 4.**
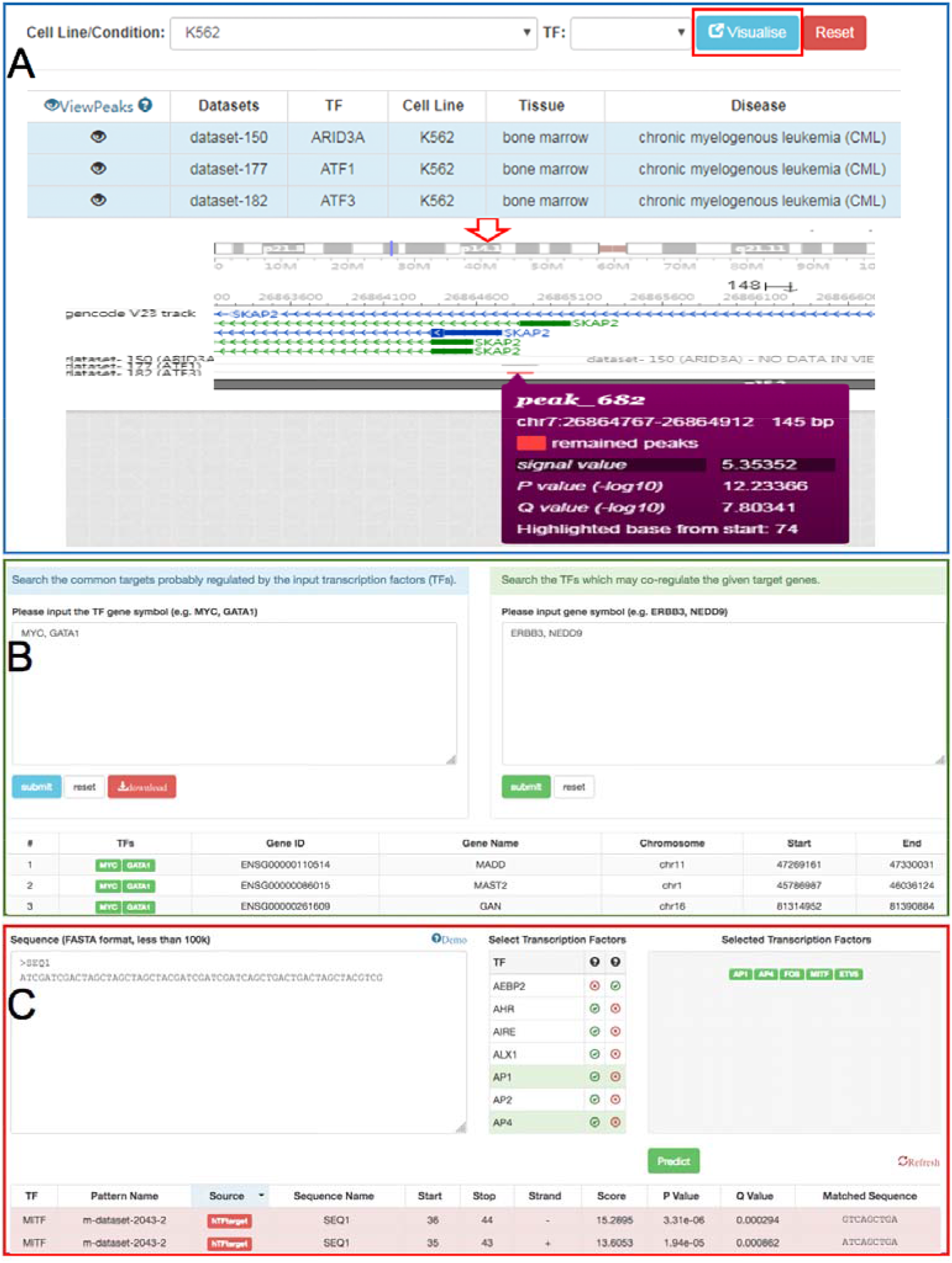
Other important modules in hTFtarget. (A) Peak visualization for TFs in user selected cell lines. (B) Potential targets and TFs co-regulation analysis for input gene sets. (C) TFBS prediction for given sequences. TFBS: transcription factor binding site.

## Discussion

Understanding TF-target regulatory relationships is a critical issue when investigating complex molecular mechanisms underlying biological processes and diseases [26,27]. Up to date, several resources have been dedicated to identify TFs, such as PlantTFDB and AnimalTFDB [28,29], and large-scale of ChIP-Seq data have been available for the detection of TF-target regulatory relationships as well. Integrative resources would offer significant insights into the dynamic TF-targets regulatory mechanisms [30]. In this study, we presented the hTFtarget, a comprehensive resource focusing on the regulation between human TFs and their targets by integrating ChIP-Seq data and TFBS prediction.

Although several resources could indirectly investigate potential regulatory interaction between TF(s) and target(s) based on ChIP-Seq peak signals, a database concisely indicating target genes for specific TFs and revealing comprehensive regulations of TF-targets is still lacking. For example, CistromeDB, collected a large number of datasets and TFs, however, it did not distinguish the TF, transcriptional co-factor, RNA polymerases, TFII family members, and their compounds. Additionally, although CistromeDB can search TF-target regulations for a single query gene and browse peak information of a given TF, it ignores the co-regulation information for TF-targets, and does not refer to the TF-target regulations at the genome-wide level. The ReMap database just provides peak information for TFs in ChIP-Seq datasets. The HOCOMOCO [22], JASPAR [23], TRANSFAC v2018 [24], and TFBSbank [31] databases focus on the collection and discovery of the motif profiles for TFBSs, while TRRUST elucidates TF–target interactions using text mining [13]. hTFtarget is the only currently available database integrating various resources to provide an almost one-step solution in investigating TF-target regulations in human.

Further development of hTFtarget will be focused on spatiotemporal classification of TF-target regulations in different gene families and diseases, such as integrating with GSCALite cancer analysis [32]. Additionally, we will continue to upgrade hTFtarget with new datasets and more epigenetic modifications, keep improving algorithms for better accuracy and offer more functions and tools. We will maintain and regularly update hTFtarget as we did for our AnimalTFDB database, which have been maintained for more than eight years. We believe that hTFtarget will be a valuable resource for the research community.

## Conclusion

The identification of TF-target regulations is an important issue for revealing the molecular mechanisms underlying complex biological processes. hTFtarget has integrated huge ChIP-Seq resources and epigenetic modification information to identify putative TF-target regulations in human. hTFtarget provides a comprehensive, reliable, and user-friendly resource, and an almost one-stop solution to explore TF-target regulations in human.

## Authors’ contributions

QZ, WL and HMZ: Data collection, bioinformatics analysis and webserver work. GYX, YRM and MX: Data collection and webserver work. QZ and WL: Conceptualization and Methodology. QZ: Conceptualization, Funding Acquisition, Manuscript writing. AYG and QZ: Conceptualization, Writing-Review and Editing, Funding Acquisition, Supervision.

## ACKNOWLEDGEMENTS

We would like to thank colleagues in groups of Ensembl, GEO, SRA, ENCODE, TRANSFAC, JASPAR, HOCOMOCO, Roadmap Epigenomics and AnimalTFDB. We also acknowledge the funding from the National Natural Science Foundation of China (NSFC) (Nos. 31822030, 31801113, and 31771458), National Key Research and Development Program of China (No. 2017YFA0700403), and China Postdoctoral Science Foundation (No. 2018M632830).

